# REM Sleep Misfires: Intruding Delta Waves Forecast Tau, Amyloid, and Forgetting in Aging

**DOI:** 10.1101/2025.08.18.670879

**Authors:** Tenzin Desel, Matthew P Walker, Chelsea Brown, Cindy Chen, Trevor A Chadwick, William J Jagust, Omer Sharon

## Abstract

Rapid Eye Movement (REM) sleep degrades with age, and more severely in Alzheimer’s disease (AD). REM sleep comprises about twenty percent of adult sleep, alternates between phasic and tonic periods, and includes delta waves (1-4Hz) in two forms: fast sawtooth waves and slower, NREM-like waves, whose expression dynamically varies across REM periods. Yet, the functional relevance of these REM sleep delta waves remains unknown. Here, using two independent cohorts, we show that aging is associated with a shift from fast sawtooth to slow NREM-like delta waves, particularly during phasic REM sleep—a period typically marked by high cortical activation. Beyond chronological age, this shift is associated with amyloid-beta and tau burden, suggesting that AD pathology disrupts REM-specific oscillatory patterns. Furthermore, this shift in REM oscillations is linked to impaired overnight memory consolidation, independent of NREM sleep quality. Moreover, variation in ApoE alleles, a major genetic risk factor for AD, was independently associated with a reduction in fast sawtooth wave density, thereby linking a genetic predisposition for AD to these specific REM microstructural changes. These findings identify a novel signature of memory decline in aging and implicate REM sleep as a distinct vulnerable substrate through which AD pathology may impair brain function.

## Introduction

REM sleep is a distinct and complex physiological state characterized by bursts of rapid eye movements (REM), wake-like asynchronous brain activity indexed by mixed-frequency EEG, profound muscle atonia, and intermittent muscle twitches^1–3^. This paradoxical^4,5^ mix of wake-like cortical activation and near-complete muscle paralysis unfurls in recurring cycles across the night. It is also non-trivial in amount, accounting for approximately twenty percent of adult sleep time^6,7^.

Although REM sleep is known for desynchronized brain activity, recent studies have revealed the presence of delta waves (1–4 Hz), typically a hallmark of NREM sleep. These include sawtooth waves (2–3Hz), which typically precede rapid eye movements,^8–11^ and slower NREM-like delta waves prominent in occipital regions^12,13^. REM delta waves are not uniformly distributed across the scalp. Instead, these REM slow waves are dominantly expressed in two topographically distinct regions^12^ 1) a frontocentral cluster of fast (2-3Hz) sawtooth waves, and 2) a medial-occipital cluster of slow NREM-like waves (<2 Hz), which are differentially modulated by phasic and tonic REM periods^12^. Phasic REM sleep, defined by bursts of REM, shows increased gamma power (35–48Hz)^14,15^, enhanced thalamocortical functional connectivity^16–18^, and reduced sensory responsiveness^14–16,18,19^ In contrast, tonic REM sleep is marked by an absence of REM, reduced cortical activation, and a weaker disconnection from external stimuli compared to phasic REM.^20–24^Importantly, delta wave expression varies across these sub-states, with phasic REM showing increased sawtooth wave density^12^.

Despite these advances, the function of REM sleep delta waves remains unknown. Futhermore,, whether such REM slow wave processes change across the lifespan, not only as a function of aging, but also in the context of abnormal aging under conditions of Alzheimer’s disease (AD) pathology (high tau and/or amyloid-beta (Aβ) burden), is similarly unknown. The latter questions are pertinent considering that REM sleep proportionally decreases with age,^25,26^ and delta wave power during REM sleep increases with age^27^. Moreover, spectral EEG slowing in REM sleep is markedly more pronounced in patients with amnestic mild cognitive impairment (aMCI) and AD, ^28,29^ where it correlates with lower cognitive scores^30^. In addition, elevated AD pathology is associated with abnormal REM sleep architecture,^31^ reinforcing a possible link between REM disruption and AD pathology.

Beyond pathological markers, genetics may also contribute to REM sleep alterations associated with AD. Apolipoprotein E (ApoE) gene, is a well-established genetic risk factor to late-onset AD ^32–36^. ApoE regulates lipid transport and exists in three polymorphic alleles ε2, ε3, and ε4, with ε4 increasing AD risk, ε3 considered neutral, and ε2 offering protection ^33^. Carriers of the ε4 allele exhibit reduced sleep efficiency, and specfically shorter REM sleep duration compared to non-carriers, further implicating ApoE-related genetic risk in sleep disruption observed in AD ^37–41^.

Building on this evidence, here, we tested the hypothesis that aging is associated with impairments in the slow delta waves of REM sleep; such changes are further exaggerated under conditions of AD-pathology load, and the extent of the age and AD-pathology-related deficits in slow oscillations, in part, statistically explain impairments in overnight memory consolidation.

## Results

In brief (and see Methods), we analyzed data from two independent cohorts who underwent overnight, in-lab polysomnographic multichannel EEG recordings. Data from the first cohort was collected at UC Berkeley and comprised cognitively unimpaired older adults (N = 70; age = 74.79 ± 5.73 years; 32% male) and younger adults (N = 55; age = 20.24 ± 1.91 years; 49% male). REM sleep epochs were identified according to American Academy of Sleep Medicine (AASM) guidelines, and rapid eye movements (REMs) were used to distinguish phasic from tonic REM periods. Building on prior work distinguishing specific subtypes of delta waves during REM sleep, oscillations were classified into sawtooth waves **(Fig. 1a**, blue dots) and slow, NREM-like waves (**Fig. 1a**, red dots) according to waveform duration (**Fig. 1b**). A subset of older participants (N=62) and younger participants (N=15) completed a sleep-dependent episodic memory test before and after overnight sleep recordings. Additionally, 59 older participants underwent PET imaging with [11C]PiB to assess Aβ levels, and 39 participants underwent [18F]Flortaucipir (FTP) to assess tau burden (**Methods**). The second cohort included 558 adults (age = 45.58 ± 13.63 years; 37.8% female) from the National Sleep Research Resource (NSRR; ^42,43^. This dataset includes overnight PSG, demographic and clinical variables, and ApoE genotype information. PSG recordings were scored in 30-second epochs using the Rechtschaffen and Kales criteria, and were preprocessed using the same pipeline as the Berkeley cohort to extract fast sawtooth and slow NREM-like wave density (**Methods**).

**Figure 1.**
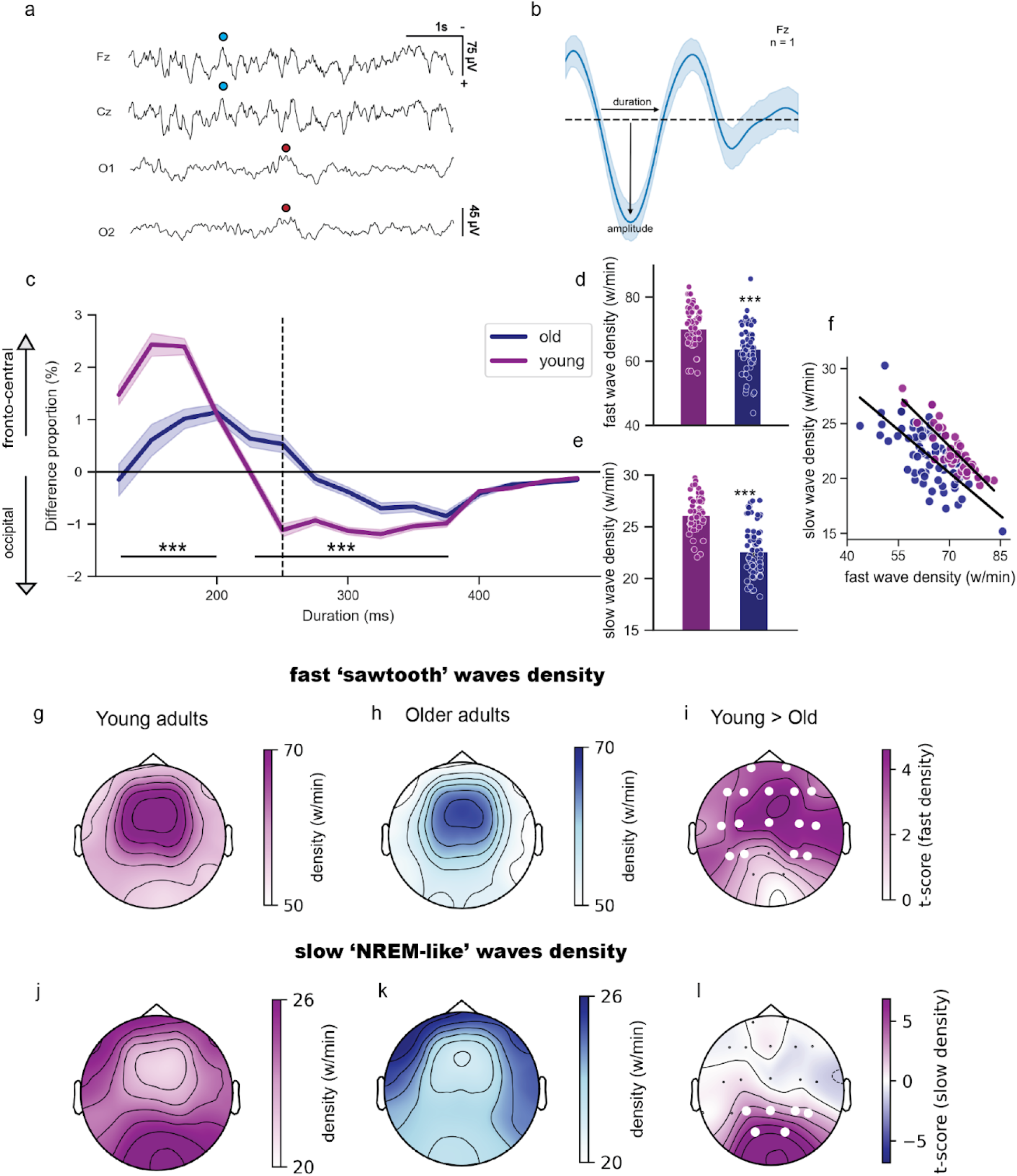
Topographic distribution of REM sleep delta waves in young and older adults. **a)** Representative raw EEG traces recorded during REM sleep in a young adult, showing fast sawtooth waves (blue dots) detected at Fz, Cz electrodes. Traces from occipital electrodes O1, O2 (below) display slow NREM-like delta waves (red dots). **b)** Averaged delta wave from Fz in the same individual, arrows indicate the waveform properties used for topographic distributions: negative half-wave duration (ms) and negative amplitude (µV) **c)** Group-level differences in the relative proportion of waves (%) across negative half-wave durations (125-500ms, in 25ms bins) between frontal (Fz) and occipital (O1/O2 averaged) electrodes in young adults (purple) and older adults (blue). Shaded areas indicate SEM. The dashed vertical line marks the 250ms threshold used to differentiate fast sawtooth waves from slow NREM-like waves **d)** Density of frontocentral fast sawtooth waves (125–250ms) in young adults (purple) and older adults (blue) **e)** Density of occipital slow NREM-like waves (250–300ms) in young adults (purple) and older adults (blue) **f)** Negative association between fast sawtooth waves and slow NREM-like waves at electrode Fz in both young (blue) and older adults (purple) **g)** Topographic maps of fast sawtooth wave density in young adults and (h) in older adults **i)** Topographic map of the t-score differences in fast sawtooth wave density between age groups; purple indicates young>old. White dots denote electrodes in the significant cluster. **j-k)** Topographic map of slow NREM-like wave density in young adults (j) and older adults (k) **l)** Topography of the t-score comparing slow NREM-like wave density across age groups; purple indicates young>old. White dots indicate electrodes in the significant cluster.

### Altered Topography of REM Delta Waves in Aging

First, we replicated prior findings in young adults^12^, confirming that REM sleep exhibits frontocentral sawtooth waves and occipital NREM-like slow delta waves. Indeed, two distinct clusters were observed: a frontocentral cluster with faster (**Fig. S1a**), denser (**Fig. S1b**), and higher amplitude waves (**Fig. S1c**), and an occipital cluster with slower (**Fig. S1d**), sparser (**Fig. S1e**), and lower amplitude waves (**Fig. S1f**).

Next, REM delta waves in older adults were analyzed using identical algorithms (**Methods**). To quantify spatial differences in wave density, the relative proportions of delta waves across frontal versus occipital electrodes were calculated^12^. Older adults showed lower frontal fast sawtooth wave proportions at 125–200 ms (t>4.0, p<0.001) and lower occipital slow NREM-like wave proportions at 250–375 ms (t>2.5, p<0.01, both FDR corrected across time bins, **Fig. 1c)**.

The morphology of **fast sawtooth waves** in older adults was similar to that of young adults - oscillating at a comparable frequency of 2.63±0.05 Hz for an average half-wave duration of 184.78±2.81ms (in young adults: 2.71±0.05 Hz, 185.25±1.91ms, respectively, p>0.05). However, the negative peak amplitude of fast sawtooth waves was reduced in older adults (7.14±1.77µV) compared to young adults (9.34±162µV, t=7.31, p=2.44×10^−11^). Similarly, **NREM-like slow waves** had comparable durations across groups (317.41±7.36 ms in older vs. 313.99±6.02ms in younger adults) and similar frequency (1.72Hz; all p>0.05). Yet, their amplitude was also significantly reduced in older adults (6.01±1.86µV) compared to young adults (8.47±2.29µV, t=6.75, p=4.56×10^−10^).

Despite similar waveform morphology, older adults exhibited a marked reduction in the density and spatial distribution of REM delta waves. Older adults showed fewer fast sawtooth waves per minute (wpm) (63.71±6.92, over Fz) compared to younger adults (70.11±9.6, t=-4.39, p<10^−4^), across a wide array of frontocentral electrodes (cluster test, p=0.001, cohen’s d=0.73, **Fig. 1h**). In occipital electrodes, older adults showed 22.56±2.65 slow NREM-like wpm, significantly less than the 26.07±2.99wpm observed in young individuals (t=-7.06, p=9.37×10^−11^). Notably, while younger adults exhibited the typical occipital cluster (Fig. 1i), it was absent in older adults (Fig. 1j; cluster test, p=0.01, cohen’s d=0.82, **Fig. 1k**). Beyond lower density, older adults exhibited a significantly reduced proportion of sawtooth waves occurring as part of a burst.^8,11,12,15^ (cluster test; p=0.03, cohen’s d=0.62), suggesting that beyond wave density, aging disrupts temporal organization (**Fig. S2**).

Next, the relationship between the two delta wave subtypes was examined. Fast sawtooth wave density was strongly negatively correlated with slow NREM-like wave density during REM sleep in young (r=-0.87, p=2.91×10^−19^; **Fig. 1f**) and in old adults (r=-0.70, p=6.46×10^−12^; **Fig. 1f**), indicating that slow NREM-like waves occur at the expense of fast sawtooth waves.

### Aging Disrupts Phasic–Tonic Modulation of REM Delta Waves

Previous work in young adults has shown that the density of delta waves is modulated across the cycle of phasic and tonic REM sub-states^12^; the next analysis tested whether these dynamics differ with aging. Focusing on a representative midline electrode (Cz), a striking divergence of delta wave expression emerged. Fast sawtooth waves density increased during phasic REM compared to tonic REM (mean change=6.39±9.92%, min=-17.17%, max=30%, t=4.66, p=2.06×10^−5^, **Fig. 2c**). In contrast, slow NREM-like waves decreased during phasic REM (-6.44±12.29% [min=-29.41%, max=22.62%], t=-4.35, p=5.99×10^−5^, **Fig. 2d**), in line with previous studies in young adults. At the scalp level, fast sawtooth waves showed a widespread increase during phasic REM relative to tonic REM (cluster test; p=0.001, cohen’s d=0.73), while slow NREM-like waves exhibited a broad suppression across the cortex during phasic REM (cluster test; p=0.001, cohen’s d=0.87, **Fig. 2j**).

**Figure 2.**
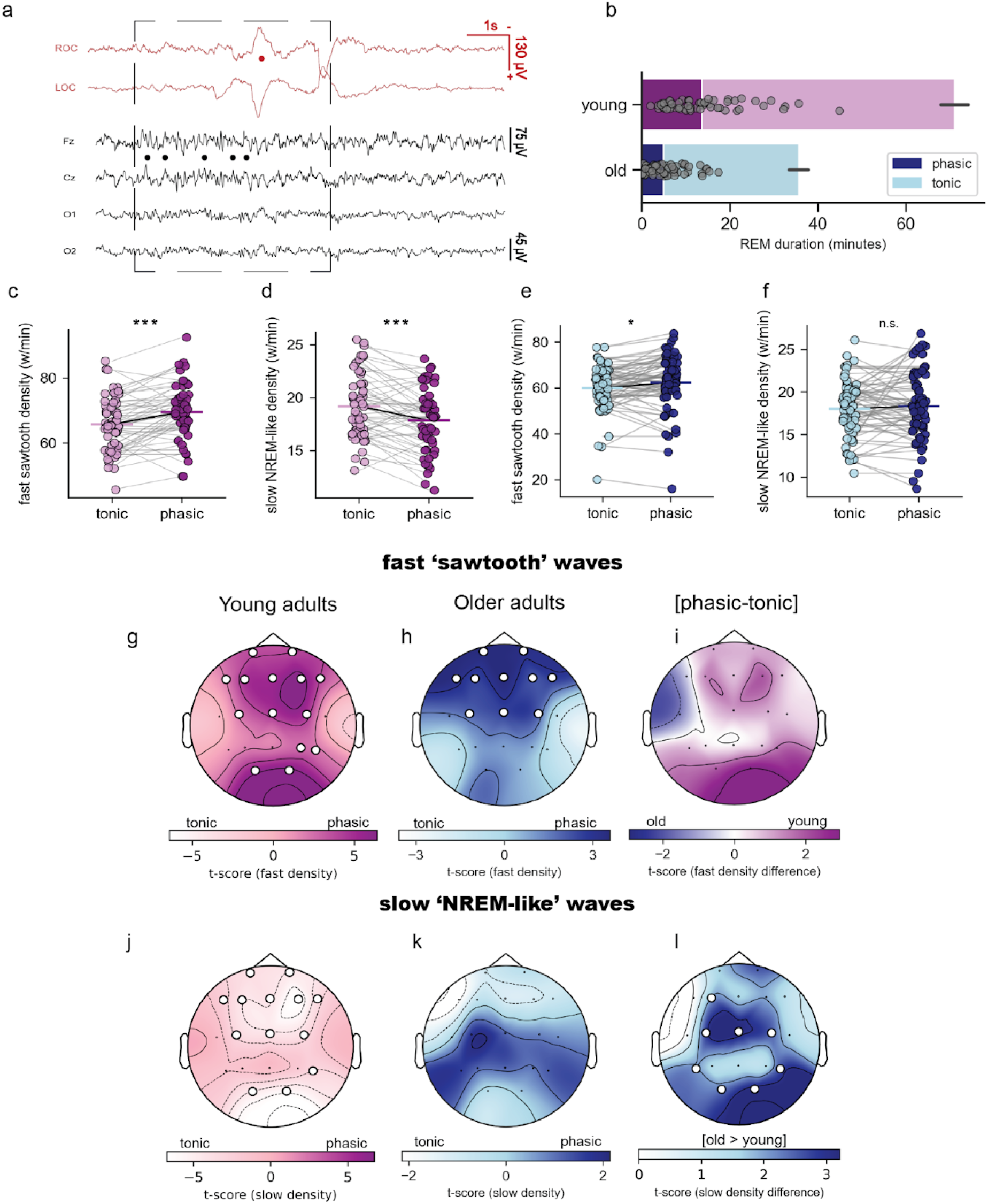
Persistence of slow NREM-like waves during phasic REM period in older adults. **a)** Representative EEG and EOG traces recorded in a young adult, showing REM (EOG, red) from the right and left outer canthi (ROC, LOC) and EEG activity (black) from the Fz, Cz, O1, and O2 electrodes. Red dots denote detected rapid eye movement (REM), and black dots indicate detected fast sawtooth waves **b)** Duration of tonic and phasic REM sleep (minutes) relative to total REM sleep duration in young adults (light pink and purple) and older adults (light blue and dark blue). Gray dots represent individual participants for the phasic REM period; error bars indicate SEM for total REM duration. **c-d)** Density of fast sawtooth waves (c) and slow NREM-like waves (d) in tonic and phasic REM sleep in young adults (Cz) with gray lines representing individual participants. **e-f)** Density of fast sawtooth waves (e) and slow NREM-like waves (f) during tonic and phasic REM sleep in older adults (Cz) with gray lines representing individual subjects. **g-h)** Topography of fast sawtooth wave density in young adults (g) and older adults (h), with white dots indicating electrodes in the significant cluster (p=0.001 for young adults; p=0.01 for older adults). **i)** Cluster permutation results comparing phasic-tonic fast sawtooth wave density difference between groups (no significant clusters). **j-k)** Topography of slow NREM-like wave density in young adults (j) and older adults (k). White dots indicate electrodes in the significant cluster (p=0.001) in young adults; no significant clusters in older adults. **l)** Cluster permutation comparing the phasic-tonic group difference in slow NREM-like wave density. White dots indicate electrodes in the significant cluster (p=0.01).

In older adults, a different pattern emerged. Fast sawtooth wave density still increased during phasic REM (+4.11±14.57%, min=-26.02, max=55.17%, t=2.51, p=0.01, **Fig. 2e**), but the density of slow NREM-like waves did not differ between phasic and tonic periods (t=0.77, p=0.43, **Fig. 2f**). Topographically, the phasic REM increase in fast sawtooth among older adults was confined to the frontocentral regions (cluster test; p=0.009, cohen’s d=0.37, **Fig. 2h**), unlike the widespread distribution seen in younger adults. For slow NREM-like waves, no significant phasic-tonic difference was observed across the scalp in older adults (cluster test; p=0.40; **Fig. 2k**). When directly comparing age groups in phasic-tonic modulation of sawtooth waves density, no group difference was observed (cluster test; p=0.16, **Fig. 2i**), suggesting that the phasic enhancement of fast sawtooth waves is preserved with age. In contrast, older adults exhibited an increase in slow NREM-like wave density during phasic REM, particularly in central electrodes (cluster test; p=0.003, cohen’s d=0.78, **Fig. 2l**), suggesting that aging entails an impairment in suppressing these slow waves and maintaining an activated EEG during REM sleep.

### A Shift in REM Sleep Delta Wave Expression Is Associated with Impaired Memory Consolidation in Aging

The decline in sleep-dependent memory consolidation with aging is well documented^44–46^. The next analysis tested whether these changes are linked to REM sleep physiology. Participants completed a word-nonword associative memory task before and after the overnight sleep recording (**Methods**). Analysis showed that a higher density of slow NREM-like delta waves in frontal and central brain regions during REM sleep was strongly associated with worse overnight memory consolidation (cluster-corrected correlation, p=0.005 **Fig. 3a**, r=-0.38, p=0.002; rho=-0.31, p=0.01, slope averaged across significant electrodes in cluster **Fig. 3b**). This association remained robust after controlling for age and sex (partial r=0.385, p = 0.002, partial rho=-0.31, p=0.01) and persisted after excluding outliers (r=-0.24; rho=-0.25; p<0.05, **Fig. 3b**). These findings suggest that slow NREM-like delta waves during REM sleep are maladaptive, particularly in frontal and central regions (**Fig. 3a**).

**Figure 3.**
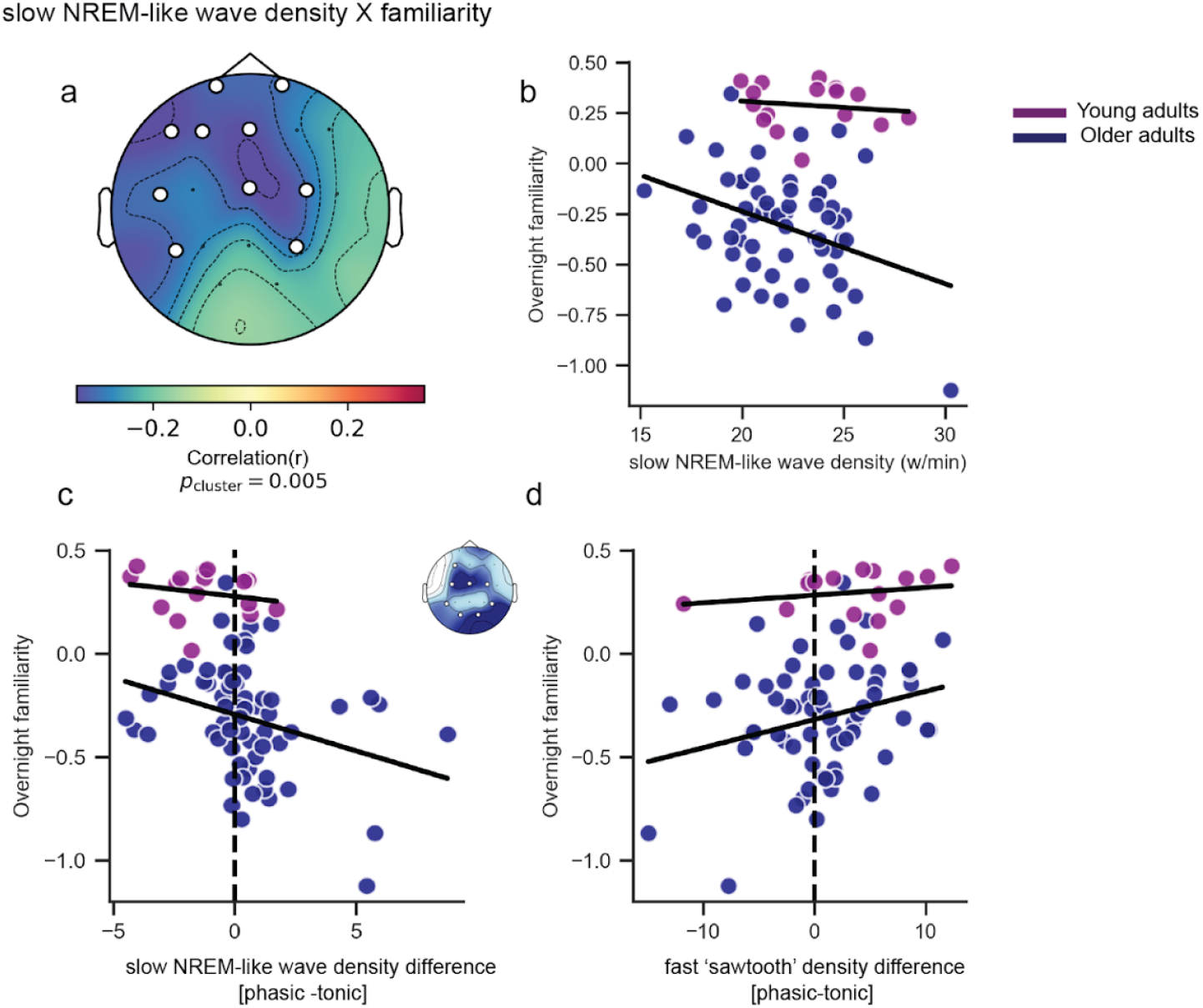
Association Between REM Sleep delta waves and memory performance. **a)** The topography of the negative correlation between slow NREM-like wave density and overnight familiarity across all REM periods using a nonparametric permutation test. The color scale represents permuted correlation r-values; White dots indicate electrodes in the significant cluster. **b)** Negative association between slow NREM-like wave density (cluster from panel a) and overnight familiarity for both young (purple) and older adults (blue). **c)** Negative association between the phasic-tonic difference in slow NREM-like wave density (waves/min; cluster shown in inset) and overnight familiarity. **d)** Positive association between the phasic-tonic difference in fast sawtooth wave density (cluster in inset) and overnight familiarity.

Subsequently, whether the reduced phasic–tonic delta waves modulation observed in older adults (**Fig. 2**) was functionally relevant was tested. In older adults, higher slow NREM-like wave density in central and occipital regions during phasic REM was significantly associated with worse overnight memory performance (r=-0.30, p=0.01; rho=-0.35, p=0.004, **Fig. 3d**, adjusted for age and sex: partial r=-0.30, p=0.01, partial rho=-0.37, p=0.003, and after excluding outliers r=-0.33, rho=-0.40, p<0.05). Conversely, fast sawtooth wave density in these same regions was associated with better overnight memory consolidation (partial r=0.27, p=0.03 **Fig. 3e**). Together, these correlations suggest that the persistence of slow NREM-like delta waves during phasic REM, at the expense of fast sawtooth waves, reflects a maladaptive shift that contributes to deficits in overnight memory consolidation.

### REM Slow Delta Waves Predict Memory Impairment Independent of NREM Sleep

Fast sawtooth waves are REM sleep specific, whereas slow NREM-like delta waves occur at comparable densities in both REM and NREM sleep^12^. Consistent with prior reports, young adults showed a higher density of fast sawtooth waves in REM compared to NREM sleep (REM (Fz): 70.11±9.61wpm, NREM: 55.04±5.50wpm, t=11.24, p=6.99×10^−6^). This preferential occurrence was preserved in older adults: REM=63.71±6.92wpm, NREM=56.44±5.98wpm, (t=8.67, p=9.54×10^−13^, **Fig. S3a**). However, older adults exhibited a higher density of sawtooth waves during NREM sleep compared to young adults (t=6.10, p=4.46×10^−9^, **Fig. S3a**), suggesting that the REM-specificity of these waves diminishes with age, mirroring the reduced selectivity also observed for slow NREM-like waves. Importantly,, frontocentral delta wave density during NREM sleep was not associated with overnight forgetting (r=-0.23, p=0.06; rho=-0.13, p=0.28, **Fig. S3c-d**). When using both NREM and REM sleep slow delta waves to predict overnight memory consolidation, only REM waves are predictive (partial r= -0.28, p=0.02 adjusted for age and sex), indicating the detrimental relationship between slow delta waves and memory consolidation is specific to REM sleep.

Next, the hypothesis that slow NREM-like waves during REM in older adults reflect a compensatory response to reduced NREM slow wave activity was tested. Classical NREM slow waves (<1.5Hz) were detected using established algorithms^47,48^ (**Methods**). Consistent with prior studies, NREM slow wave density (partial r=0.32, p=0.01) and cortical involvement (partial r=0.33, p=0.01) predicted better overnight memory consolidation. However, REM slow delta waves density remained a significant negative predictor of retention after accounting for NREM slow wave density and their cortical involvement (partial r=-0.32, p=0.01; **Table. S4 and S5**). Moreover, REM slow delta wave density was not correlated to NREM slow wave density (r=-0.07, p=0.55), arguing against a compensatory relationship. These results suggest that excessive REM slow waves exert an additional, independent, REM-specific detrimental effect on memory consolidation.

### Alzheimer’s Disease Pathology Is Associated with Altered Phasic–Tonic Delta Wave Composition

Having established that persistent slow NREM-like waves during phasic REM are maladaptive for overnight memory consolidation, we next investigated whether variability in delta wave dynamics among older adults was related to Alzheimer’s disease (AD) pathology, specifically Aβ (PiB-PET) and tau (FTP-PET).

Older adults classified as Aβ-positive (Aβ PiB>1.065) exhibited a significantly higher density of slow NREM-like waves during the phasic REM period compared to Aβ-negative older adults, particularly over central electrodes (**Fig. 4a**, cluster test; p=0.045, cohen’s d=0.63; t=2.95, **Fig. 4b**, p=0.004). While Aβ-negative older adults exhibited a significant suppression of slow NREM-like wave activity during phasic REM, primarily in frontal regions (cluster test; p=0.01, cohen’s d=0.65, **Fig. 4c**), similar to young adults (**Fig. 2j**), Aβ-positive individuals showed no significant difference (cluster test; p=0.07, **Fig. 4d**).

**Figure 4.**
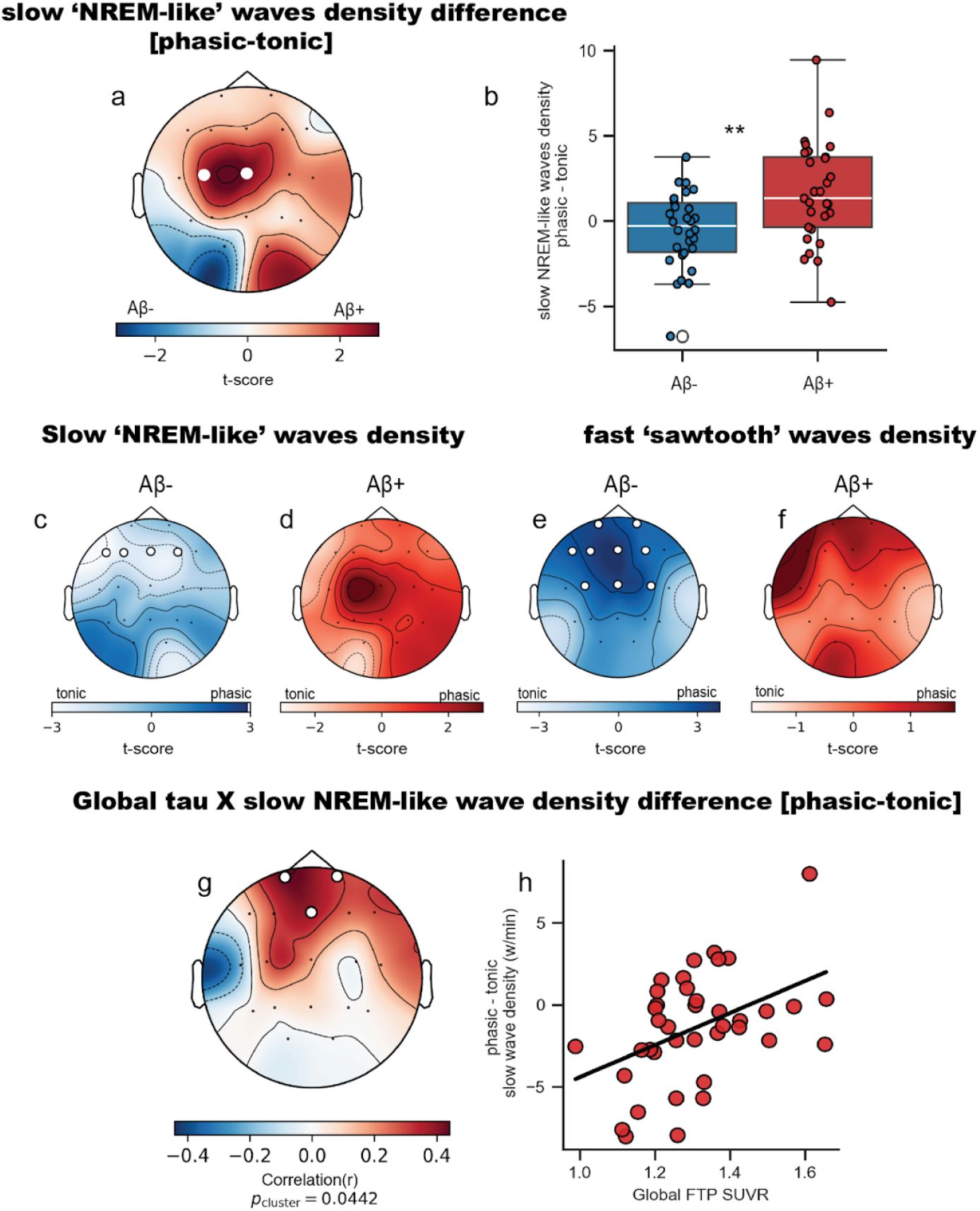
Greater persistence of slow NREM-like waves during the phasic period in Aβ+ older adults, and high tau burden, predicts greater persistence. Topography of the phasic-tonic difference in slow NREM-like wave density between Aβ-negative and Aβ-positive older adults (cluster permutations). White dots indicate electrodes within the significant cluster. **b)** phasic-tonic difference in slow NREM-like wave density in Aβ-negative and Aβ-positive older adults, averaged over the significant cluster from panel a. **c-d)** Topography of slow NREM-like wave density in Aβ-negative (c) and Aβ-positive (d) older adults. White dots indicate electrodes in the significant cluster in Aβ-negative individuals; no significant cluster in Aβ-positive individuals. **(e-f)** Topography of fast sawtooth wave density in Aβ-negative (e) and Aβ-positive (f) older adults. White dots indicate electrodes in the significant cluster in Aβ-negative individuals; no significant cluster in Aβ-positive individuals. Topography of slow NREM-like wave density in Aβ-positive individuals (no significant cluster**). (g)** Non-parametric permutation analysis shows a positive correlation between global tau burden (FTP SUVR) and the phasic-tonic difference in slow NREM-like wave density (waves/min). White dots indicate electrodes within the significant cluster. **(h)** Positive association between global tau levels and the phasic-tonic difference in slow NREM-like wave density, averaged over the significant cluster identified in panel g.

For fast sawtooth waves, the density difference between phasic and tonic periods did not differ between Aβ-negative and Aβ-positive older adults (cluster test; p>0.05). However, Aβ-negative older adults show an increase in fast sawtooth wave during phasic REM, particularly over frontocentral regions (cluster test; p=0.006, d=0.57, **Fig. 4e**). This effect was absent in Aβ-positive older adults (no significant clusters, **Fig. 4f**).

To further investigate the influence of tau pathology on the persistence of slow NREM-like waves during phasic REM sleep was assessed. Global tau burden was predictive of slow NREM-like wave activity in frontal regions (cluster correlation; p=0.04, **Fig. 4g**, significant also after adjusting for age and sex; partial r=0.40, p=0.01, **Fig. 4h**). To identify regional specificity, delta wave density differences within the significant cluster were extracted and correlated with tau burden across all cortical regions in the Desikan-Killiany Atlas (DKT). Higher tau burden in the precentral, postcentral, superior parietal, caudal middle frontal, superior temporal, inferior temporal, and fusiform regions (FDR corrected) was significantly predictive of greater slow NREM-like wave density in the phasic period, after adjusting for age, sex, global Aβ burden, and cortical thickness, and FDR correction for multiple comparison (**Fig. S6a**). In sum, both Aβ and tau accumulation are associated with the maladaptive persistence of frontocentral slow NREM-like waves during phasic REM sleep.

Finally, the specificity of pathological REM sleep EEG slowing was tested by comparing identically detected NREM-like delta waves in REM sleep and wakefulness. Slow NREM-like wave density was higher during REM sleep (cluster test, p=0.001, **Fig. S4.a**). Moreover, Aβ-positive and Aβ-negative individuals showed no difference in wakefulness slow delta waves density (no significant cluster, **Fig. S4.b**), and slow waves during wakefulness showed no association with overnight memory consolidation (r=-0.02, p=0.87, **Fig. S4.c**). This suggests that the increase in slow NREM-like waves, associated with AD pathology and memory impairment, is specific to REM sleep, not wakefulness.

### Independent Cohort Validates Age-Dependent Reduction and APOE ϵ4-Related Deficits in Phasic Fast Sawtooth Waves

Next, to assess the replicability and generalizability of the findings, delta waves were detected in an independent cohort using the same methods (n = 558, mean age = 45.58±13.63, 37.8% females, see **Methods** for additional details). In accordance with the Berkeley cohort, older age was associated with lower phasic fast sawtooth waves density (rho=–0.13, p=0.001, all channels averaged; **Fig. 5a**) Additionally, older age was associated higher rate of slow NREM-like wave density in during the phasic REM period, relative to tonic (rho=0.10, p=0.009, all channels averaged; **Fig. 5b**). Given that both amyloid and tau were linked to increased phasic slow wave intrusion, the next analysis examined whether REM sleep delta waves are related to variation in ApoE ε4 allele, a genetic risk factor for Alzheimer’s disease.

**Figure 5.**
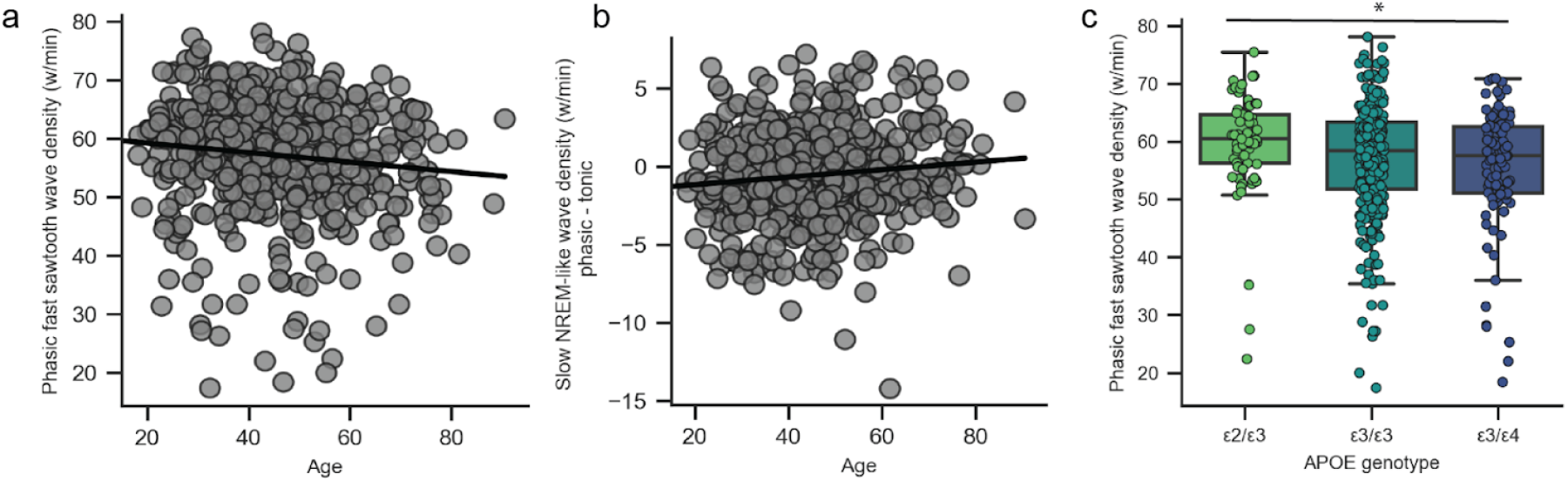
Aging increases phasic slow NREM-like activity, ApoE ε4 carriers have lower phasic sawtooth waves. **a)** Negative association between age and phasic fast sawtooth wave density (all channels averaged) **b)** Positive association between age and phasic-tonic difference in slow NREM-like wave density **c)** Phasic REM fast sawtooth wave density (averaged across all EEG channels) across ApoE genotypes. A significant reduction in phasic sawtooth wave density was observed in ε3/ε4 individuals (increased AD risk) compared to ε2/ε3 individuals (lower AD risk).

Individuals with the ε3/ε4 genotype (higher risk) showed reduced phasic fast sawtooth wave density compared to those with the ε2/ε3 genotype (lower risk) (Kruskal-Wallis H=7.633, p= 0.02; Dunn’s post hoc test, p=0.02 (**Fig. 5c**), based on an analysis restricted to the common alleles combinactions: ε2/ε3, ε3/ε3, and ε3/ε4 groups (N=490).There were no differences between ApoE genetic groups in age, REM sleep percentage, or sleep efficiency (all ps > 0.40, **Fig. S7**), and no interaction between fast sawtooth waves density,ApoE status, and age (all ps > 0.18).Moreover, even when limiting the analysis to participants younger than 50, ε3/ε4 carriers continued to show significantly lower phasic sawtooth wave density (H = 7.256, p = 0.02; Dunn’s post hoc test, p=0.02), suggesting that lower sawtooth density in ApoEε4 carriers is expressed early in life and decreases further with age.

## Discussion

The current findings establish a critical and previously underappreciated relationship between aging, AD pathology, and disruptions in REM sleep neurophysiology. Aging significantly reshapes the topographic and temporal organization of REM delta waves, marked by decreased fast sawtooth wave density, reduced occipital slow NREM-like delta activity, and notably, a maladaptive persistence of slow delta waves during phasic REM periods—a state of heightened cortical activation. This atypical delta intrusion into phasic REM was associated with impaired overnight memory consolidation and linked to elevated Aβ and tau pathology, thus identifying a novel electrophysiological marker predictive of cognitive decline. In an independent cohort, variation in ApoEε4 alleles carrier—a major genetic risk factor for Alzheimer’s Disease—was independently associated with reduced fast sawtooth wave density, further linking genetic susceptibility for AD with specific microstructural alterations in REM sleep.

Although age-related impairments in NREM sleep are well documented,^49^ alterations in REM sleep beyond reduced duration in aging have remained largely unexplored. The present results demonstrate that aging disrupts not just REM sleep quantity but crucially alters REM electrophysiological microarchitecture, reinforcing REM sleep’s critical neurocognitive functions. Specifically, marked reductions in central fast sawtooth waves were observed, which normally reflect a highly active brain state characterized by widespread gamma oscillations^14,15^ and often preceded by synchronous sawtooth waves in the posterior thalamus.^18^ In contrast, the abnormal persistence of NREM-like slow waves, normally suppressed during phasic REM^12^ possibly due to heightened cholinergic activity,^50^ may destabilize the activated, phasic REM state.

Critically, the persistence of slow NREM-like waves into phasic REM correlated negatively with overnight episodic memory retention. This suggests these slow oscillations are maladaptive, disrupting the memory-enhancing role typically attributed to phasic REM periods^51,52^. Conversely, preservation of fast sawtooth waves predicted better memory outcomes, supporting their beneficial role in memory consolidation, potentially by driving localized gamma across mnemonic, premotor, and limbic areas^15^. Moreover, the link between REM delta disruptions and memory impairments persisted independently from NREM sleep deficits, emphasizing a distinct REM-specific vulnerability in aging and AD pathology. This independence underscores that the unique neurochemical and physiological conditions of REM sleep, such as minimal noradrenergic tone, maximal cholinergic tone, and muscle atonia, play a distinct role in brain plasticity.

The association between REM delta wave disruptions and Alzheimer’s pathology is of notable significance. Elevated Aβ pathology corresponded with impaired suppression of slow NREM-like delta waves in phasic REM, aligning with previous findings of disrupted cholinergic systems in early Alzheimer’s^53^. Furthermore, increased tau burden in inferior and superior temporal cortices, as well as precentral and postcentral gyri, was correlated with persistent slow wave expression during phasic REM—overlapping prior source localization^12,14^ and intracranial recordings^15^ of sawtooth waves—suggesting that tau might impair generation of fast sawtooth waves in the cortex.

These findings call for a conceptual shift. Rather than viewing REM sleep as a passive state vulnerable to degeneration, these results propose REM sleep is an actively regulated state whose physiological precision is critical for cognitive resilience against neurodegeneration, adding to that of NREM sleep.^46,54^ This may open the door to REM-focused interventions for cognitive decline, or at least raise awareness of accelerated REM sleep deterioration in aging.

Additionally, this work raises questions about the interplay between neuromodulatory imbalance and AD pathology progression. The specific vulnerability of phasic REM to tau and Aβ pathology implies a selective disruption of microcircuits responsible for high-frequency oscillations essential for memory^53,55–60^. Identifying these circuits and their vulnerability could pave the way for targeted neuromodulation therapies.

Taken together, these data support a model wherein aging and incipient Alzheimer’s pathology disrupt oscillatory dynamics during REM sleep, in part through tau-associated cortical dysfunction and impaired cholinergic modulation. ApoEε4 carriers express REM delta waves patterns associated with an aged brain compared to those with low risk ε2 allele. These disruptions not only serve as early electrophysiological biomarkers for AD risk but also associate with impaired episodic memory, the prototypical hallmark of cognitive decline.

## Methods

### Participants

Seventy cognitively normal older adults (mean age=74.79, SD±5.73, 32% male) were recruited from the Berkeley Aging Cohort Study (BACS). Fifty-five healthy young adults (mean age=20.24, SD±1.91, 49% male) were recruited from the UC Berkeley student community. Data from both groups have been included in previous publications^44,45,61,62^. Exclusion criteria included any reported psychiatric, neurological, or sleep disorders or use of hypnotic or antidepressant medications. Additionally, all participants scored above 25 in the Mini-Mental State Examination (MMSE)^63^, indicating unimpaired cognitive function. Electrooculogram (EOG) data were available for 69 older adults (with one excluded due to insufficient phasic activity) and 55 younger adults for the analysis of phasic and tonic REM periods. For the analysis of PET-derived AD pathology, 59 older adults had available PiB Aβ scans, and 39 older adults had FTP tau scans.

### Polysomnographic recordings

Overnight polysomnographic recordings were collected using a 19-channel 10-20 EEG system, with data digitized at 400Hz on a Grass Technologies Comet XL system (Astro-Med). Electrooculogram (EOG) recordings were placed at the right and left outer canthi (ROC and LOC, respectively) to monitor eye movements. Electromyography (EMG) was recorded from the chin to track muscular activity, according to the American Association of Sleep Medicine (AASM) guidelines^64^. Reference electrodes were placed on both the left and right mastoids. All participants abstained from caffeine and alcohol for 24 hours before their sleep study session and completed two overnight sleep sessions. The first session served as an adaptation and screening night, and the second session served as the experimental session, during which they performed episodic paired-association learning tasks before and after the sleep recordings.

### Episodic memory task

As previously described^46^, participants completed an episodic associative task involving word-nonword pairings before and after a full night of sleep (see ^44,61^ for details). All participants first completed a training session until they achieved 100% accuracy. They were then given two recognition memory tests: one immediately after the training session and a delayed test the following morning. During the recognition tests, participants were prompted with real words and were asked to choose from four options: learned pair, lure, false alarm, or new. Two memory scores were calculated: familiarity accuracy and associative recognition. Familiarity retention was measured by subtracting the rate of falsely recognized new pairs from correctly recognized studied pairs, while recollection was assessed by subtracting the rate of falsely paired nonword lures from correctly studied pairs. The paired associative learning task is sensitive to sleep-dependent episodic memory retention, with familiarity being especially sensitive to aging^65,66^.

### EEG data preprocessing

Sleep scoring was performed according to the AASM^64,67^. Next, the EEG was preprocessed using the MNE-Python package^68^. Visually obvious artifact-free EEG scored as REM were extracted from all subjects, bandpass filtered between 0.5-40 Hz using a finite impulse response (FIR) filter, and downsampled to 200 Hz. Independent component analysis (ICA) was then applied to remove ocular and electrocardiographic artifacts, as is standard for wake EEG^68^.

### Delta waves detection and classification

Delta wave detection followed methodologies established in a previous study in younger adults^12^ as implemented via the slow wave detection algorithm from the YASA Python package^69^. Signals from each electrode were downsampled to 128 Hz and bandpass filtered between 1 and 10 Hz to minimize wave shape and amplitude distortions. Delta waves (1–4 Hz) with negative half-wave durations between 125 ms and 500 ms were identified. Each electrode signal was further bandpass filtered between 1–4 Hz using a FIR filter with a 0.2 Hz transition band. Oscillations were detected based on zero-crossings of the EEG signal, focusing on negative half-waves within the specified duration range. The negative phase duration was determined by identifying the zero-crossings of the EEG signal. For each detected wave, the negative amplitude (μV), negative half-wave duration (ms), and wave density (waves per minute) were analyzed. To examine the temporal burst characteristics of sawtooth waves, a burst was defined as a sequence of three or more consecutive waves, each separated by less than 750 ms, with each wave’s amplitude not exceeding 25% of the preceding wave^12^.

### Rapid eye movement (REM) detection and classification of phasic and tonic REM

Individual rapid eye movements were detected using a previously established algorithm^70,71^ as implemented in the YASA Python package^69^. EOG signals were first bandpass filtered (1–5 Hz) using a fourth-order digital Butterworth filter to reduce slow eye movement contamination and high-frequency noise. Candidate REMs (CREMs) were identified based on the negative instantaneous product (NIP) of the two filtered EOG channels, ensuring phase-reversed synchrony. A peak detection method was applied within a sliding window to prevent detections from occurring too close together. Each CREM was then characterized by several features: artifact amplitude, NIP magnitude, inter-channel correlation, and deflection angle. CREMs exceeding artifact amplitude thresholds, with insufficient negative correlation, low NIP values, or slow deflection angles were rejected to optimize sensitivity and minimize false detections. Valid REMs were those meeting all criteria. Consecutive REMs separated by less than 1 second were identified as REM bursts and included in subsequent analysis. Phasic REM periods were defined as continuous REM epochs longer than 5 seconds, containing a burst of rapid eye movement. Tonic periods were defined as 5-second epochs without any detected rapid eye movements.

### NREM analysis - delta waves (1-4 Hz) and slow oscillations (0.3-1 Hz)

The preprocessing and detection pipeline applied to REM sleep epochs was similarly applied to artifact-free NREM epochs. Delta waves within the 1-4 Hz range and with half-wave durations between 125-500 ms were identified. The mean density and amplitude of both fast sawtooth waves and slow NREM-like waves were compared relative to REM sleep.

Additionally, individual slow waves were detected (slow oscillations 0.3-1.5 Hz) using the YASA python package^69^. Each electrode’s signal was bandpass filtered between 0.3-1 Hz using an FIR filter with a 0.2 Hz transition band. Negative peaks between -40 and -200 μV and positive peaks between 10 and 150 μV were identified. For each negative peak (representing the slow wave trough), the nearest positive peak was located, and the peak-to-peak amplitude was calculated. The durations of the negative and positive phases were measured: the negative phase was defined as the time between the first zero-crossing before the negative peak and the second zero-crossing between the peaks, while the positive phase was the time between the zero-crossing between the peaks and the last zero-crossing after the positive peak. Only waves with a peak-to-peak amplitude between 75 and 300 μV, a negative phase duration between 0.1 and 1.5 seconds, and a positive phase duration between 0.1-1 second were included in the analysis. To investigate cortical involvement, slow wave detections were sorted by the negative peak with 10 ms resolution, and within that range, by the amplitude of the negative peak. The first electrode to detect a valid negative peak was considered the origin, and additional electrodes detecting negative peaks before the mid-point of the previous wave were grouped as part of the same multichannel slow wave.

### PET acquisition and processing

Details on PET data acquisition have been described in prior studies^72–74^. Both flortaucipir (FTP) and Pittsburgh Compound B (PiB) PET tracers were produced at the Biomedical Isotope Facility at Lawrence Berkeley National Laboratory (LBNL), and all scans were conducted using a Biograph PET/CT scanner. Typically, FTP scans were performed on the same day as PiB scans, immediately after the PiB imaging session.

For FTP imaging, participants received an injection of 10 mCi of the tracer, and the analysis utilized data collected between 80 and 100 minutes post-injection. CT scans acquired before emission data collection were used for attenuation correction. FTP-PET images were reconstructed using an ordered subset expectation maximization algorithm with scatter correction applied and smoothed with a 4-mm Gaussian kernel. To create FTP standardized uptake value ratio (SUVR) images, tracer retention from the 80–100 minute time window was normalized to retention in the inferior cerebellar gray matter, serving as the reference region. Partial volume correction (PVC) was implemented using the Rousset geometric transfer matrix method to address atrophy-related effects and spillover signals^75,76^.

For PiB-PET imaging, participants were injected with 15 mCi of PiB tracer, and dynamic data acquisition occurred over 90 minutes, starting immediately post-injection. CT scans performed before tracer administration were used for attenuation correction. PiB-PET images were reconstructed using the same ordered subset expectation maximization algorithm with scatter correction and smoothed with a 4-mm Gaussian kernel. Distribution volume ratio (DVR) values were computed using Logan graphical analysis^77,78^ on data from the 35–90 minute post-injection interval, normalized to the whole cerebellar gray matter as the reference region. Global PiB metrics were derived by aggregating data from multiple cortical regions of interest identified through FreeSurfer, as outlined in prior work^79^. A standard DVR cutoff of 1.065 was used to categorize individuals as Aβ-positive or Aβ-negative^80,81^.

### ApoE independent NSRR dataset

Data from the publicly available ApoE dataset (n = 712), provided by the National Sleep Research Resource (NSRR) were analyzed, which was originally collected as part of an NIH-supported study at Stanford University between 2003 and 2007 that investigated genetic associations and clinical features in ApoE ε4 positive and negative subjects with various degrees of sleep apnea^42,43^. Participants were eligible if they had suspected but untreated sleep-disordered breathing; individuals using continuous positive airway pressure (CPAP) were excluded. This dataset includes overnight PSG with a total of 18 recorded signals—covering EEG, EOG, ECG, combined leg and submentalis EMG, snoring, respiratory effort, airflow, oxygen saturation—along with demographic, clinical, lipid profile, and ApoE genotype data. PSG recordings were acquired using Sandman Elite software with SD32+ amplifiers and manually scored in 30-second epochs according to the Rechtschaffen and Kales criteria. ApoE ε4 status was determined from blood samples and used to stratify participants into carrier and non-carrier groups into six ApoE genotype groups: ε2/ε2, ε2/ε3, ε3/ε3, ε2/ε4, ε3/ε4, and ε4/ε4.

We applied the same EEG and EOG preprocessing pipeline as described in our primary analysis. REM sleep epochs were extracted and preprocessed (n = 624); EEG signals were bandpass filtered between 0.5–40 Hz, and data were downsampled to 200 Hz. Independent component analysis (ICA) was performed using MNE-Python to remove ocular and cardiac artifacts. Delta waves were detected using the YASA Python package following previously established criteria. EEG signals were bandpass filtered between 1–4 Hz and downsampled to 128 Hz. Negative half-waves with durations of 125–500 ms were identified, and wave density (waves per minute) was calculated at each electrode. Rapid eye movements (REMs) were detected from bandpass-filtered EOG signals using a previously validated algorithm. REM bursts and phasic/tonic REM periods were defined as in our primary analysis. All preprocessing and analyses were performed using MNE-Python and YASA to maintain consistency with our original pipeline. Phasic and tonic periods were then extracted based on EOG activity, and recordings without sufficient phasic REM were excluded, yielding a final sample of 558 participants (n = 558, age = 45.58 ±13.63, 37.8% female).

### Experimental design and statistical analysis

In statistical analyses, group and conditions comparisons were conducted using independent and paired t-tests, respectively utilizing the Scipy^82^ Python package^82^. Pearson’s correlation coefficients were calculated to examine associations between wave density and behavioral performance unless specified otherwise. Partial correlations were reported after adjusting for confounding variables ^83^. For comparisons involving three or more groups with non-normally distributed data, Kruskal–Wallis tests were applied, followed by post hoc Dunn’s test when appropriate.

To assess the robustness of the correlation results (Fig.3a), a cluster-based non-parametric permutation test was performed using the Eelbrain Python package^84^, designed for EEG and MEG data analysis^84^. This approach identifies clusters of significant correlations while controlling for multiple comparisons. The strength of correlations between REM delta density and behavioral measures was further examined using the Robust Correlation Toolbox^85^ which computes correlation on data cleaned for bivariate outliers identified via the mid-covariance determinant and removed using orthogonal distances and the interquartile range threshold.

For multiple comparison corrections, MNE-python^68^ cluster-based permutation tests were employed^68^. Clusters were identified spatially by applying a two-tailed, paired and independent t-test threshold at p < 0.05, with permutations generated by randomly shuffling condition labels or groups. Clusters with a p-value of less than 0.05 (two-sided) were considered significant.

## Acknowledgements

We thank the participants for their ongoing commitment to the study. This work was supported by the National Institute on Aging, National Institutes of Health, grant numbers RF1AG054106 awarded to M.P.W. The content is solely the responsibility of the authors and does not necessarily represent the official views of the National Institutes of Health. This research was conducted while O.S. was a Glenn Foundation for Medical Research Postdoctoral Fellow.

The Sleep Disordered Breathing, ApoE and Lipid Metabolism Study was supported by the National Institute of Health (5R01HL71515-3). The National Sleep Research Resource was supported by the U.S. National Institutes of Health, National Heart Lung and Blood Institute (R24 HL114473, 75N92019R002).

