## supplementary material for "REM Sleep Misfires: Intruding Delta Waves Forecast Tau, Amyloid, and Forgetting in Aging"

### Supplementary Materials

**Figure S1 A. Topography of REM delta waves in young (replication) and older adults**

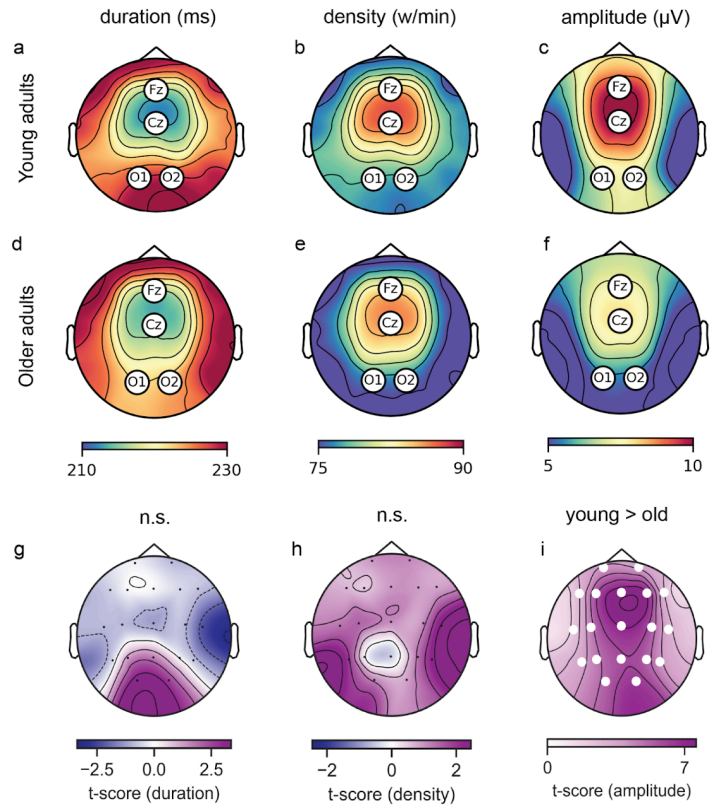

**Figure S1.** From left to right, the plots show **a)** the average density of delta waves (waves/min), **b)** negative half-wave duration (ms), and **c)** the average negative amplitude ($\mu V$) for young adults. **d-f)** The same wave properties for older adults. **g)** cluster permutation analysis comparing negative half-wave duration of delta waves between older adults (blue) and young adults (purple) (no significant cluster). **h)** cluster permutation analysis comparing the density of delta waves between groups (no significant cluster). **i)** cluster permutation analysis comparing negative amplitude of delta waves between groups (cluster test; $p = 0.001$, purple indicates young > old).

### Figure S2. Temporal sawtooth burst characteristics

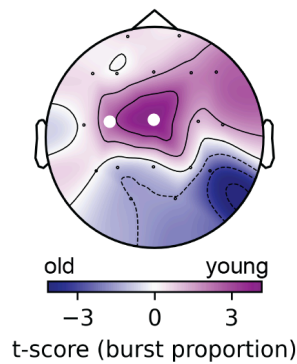

**Figure S2.** Topography of cluster permutation analysis comparing the proportion of sawtooth waves occurring in bursts between young adults and old adults, white electrodes represent a significant cluster (cluster test, $p = 0.03$).

### Figure S3. Fast sawtooth waves and slow NREM-like waves and their association with memory performance in NREM Sleep

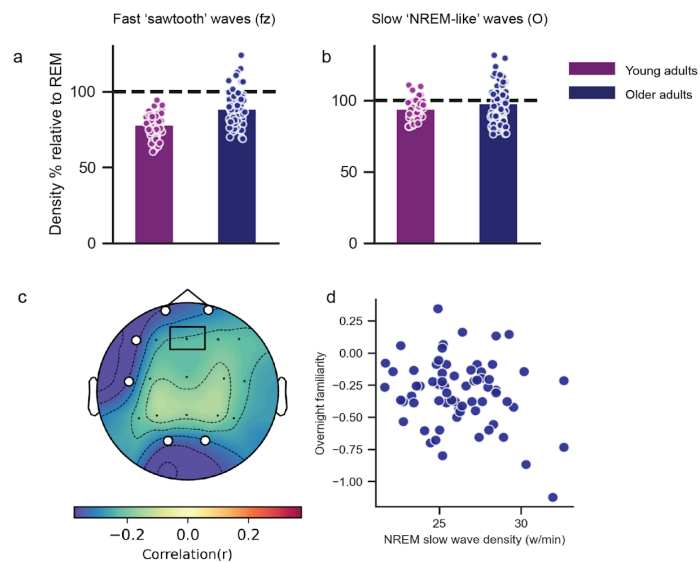

**Figure S3.** **a)** Frontocentral fast sawtooth wave density in NREM sleep relative to REM sleep. **b)** Occipital slow NREM-like wave density in NREM sleep relative to REM sleep. **c)** A non-parametric permutation correlation between slow wave density in NREM sleep and overnight familiarity. Color bars represent permuted correlation values. The black rectangle on the topography highlights the Fz channel used for the scatterplot below. **d)** No association between slow NREM-like wave density in NREM sleep at Fz electrode and overnight familiarity in older adults.

**Figure S4. Increased slow NREM-like waves in REM sleep compared to wake periods**

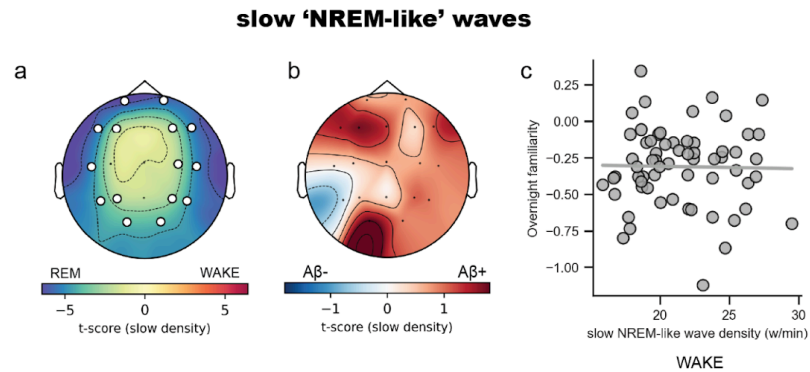

**Figure S4. a)** Topography of the cluster permutation analysis comparing the density of slow NREM-like waves between REM sleep and wake periods in all older adults, white electrodes represent significant clusters (cluster test, $p = 0.001$). **b)** Topography of the cluster permutation analysis comparing the density of slow NREM-like waves in wake periods between Aβ-negative and Aβ-positive older adults (no significant cluster). **c)** No association between slow NREM-like wave density at Fz electrode in wake periods with overnight familiarity in older adults.

**Table S4. Linear regression of NREM SO cortical involvement and REM slow delta density**

|  | coefficient | standard error | t-value | p-value | CI[2.5%] | CI[97.5%] |
| --- | --- | --- | --- | --- | --- | --- |
| Intercept | -0.288 | 0.617 | -0.467 | 0.642 | -1.525 | 0.948 |
| REM slow delta density | -0.031 | 0.012 | -2.612 | 0.012 | -0.056 | -0.007 |
| NREM SO cortical involvement | 0.033 | 0.014 | 2.428 | 0.018 | 0.006 | 0.06 |
| Age | 0.003 | 0.006 | 0.561 | 0.577 | -0.009 | 0.016 |
| Gender | -0.02 | 0.07 | -0.314 | 0.755 | -0.162 | 0.118 |

**Table S5. Linear regression of NREM SO density and REM slow delta density**

|  | coefficient | standard error | t-value | p-value | CI[2.5%] | CI[97.5%] |
| --- | --- | --- | --- | --- | --- | --- |
| Intercept | 0.177 | 0.595 | 0.297 | 0.768 | -1.019 | 1.373 |
| NREM SO density | 0.065 | 0.029 | 2.208 | 0.032 | 0.006 | 0.123 |
| REM slow delta density | -0.03 | 0.013 | -2.328 | 0.024 | -0.056 | -0.004 |
| Age | 0.001 | 0.007 | 0.191 | 0.849 | -0.012 | 0.015 |
| Gender | -0.06 | 0.08 | -0.75 | 0.457 | -0.221 | 0.101 |

**Figure S6. Whole brain spatial map of tau burden and greater persistence of slow NREM-like waves in phasic REM period**

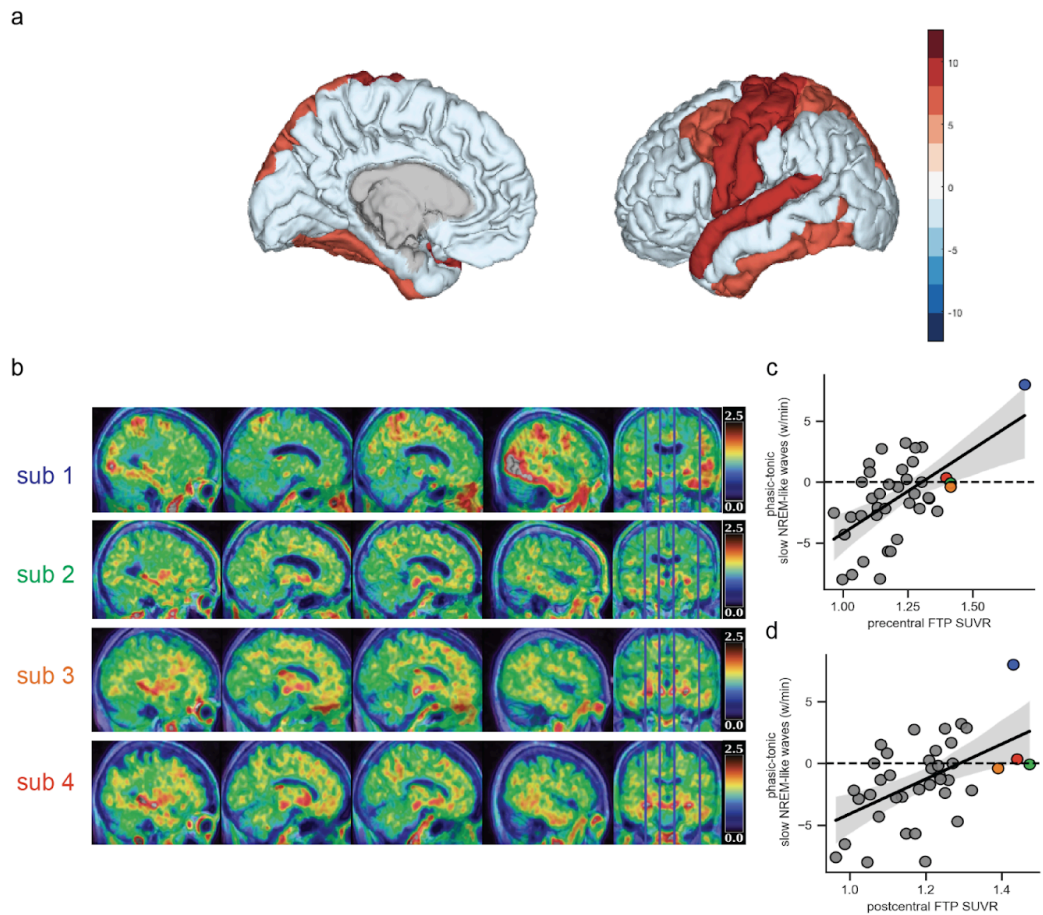

**Fig S6. a)** Tau burden in the precentral, postcentral, superior parietal, caudal middle frontal, superior temporal, inferior temporal, and fusiform regions predict greater slow NREM-like wave density in the phasic period; color bars represent beta values adjusted for age, sex, PiB, and atrophy. **b)** Tau-PET images of four individual subjects with high tau-PET signal in precentral and postcentral ROI. **c)** Positive association between precentral FTP SUVR and phasic-tonic difference in slow NREM-like waves density (subjects from panel b are highlighted accordingly). **d)** Positive association between postcentral FTP SUVR and phasic-tonic difference in slow NREM-like waves density (subjects from panel b are highlighted accordingly).

**Figure S7. No difference in age, REM sleep % and sleep efficiency in ApoE groups**

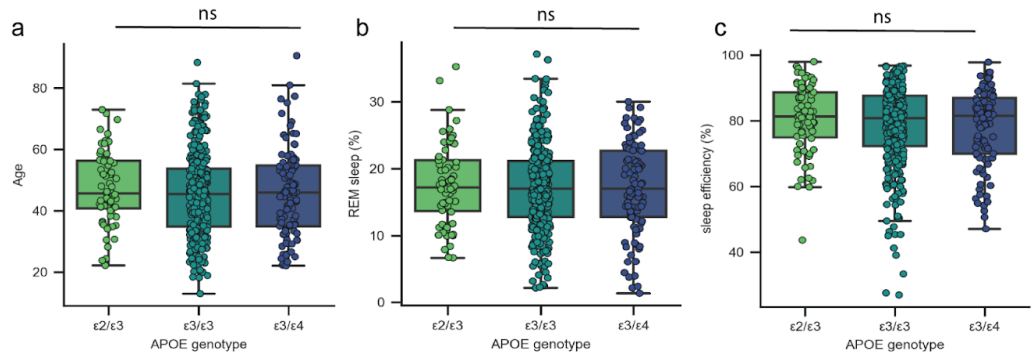

**Fig. S7.** Age (a), REM sleep percentage (b), and sleep efficiency (c) across ApoE genotype groups; no significant group differences observed in any measure.
